# Global Reduction of Information Exchange during Anesthetic-Induced Unconsciousness

**DOI:** 10.1101/074658

**Authors:** Christina Hamilton, Yuncong Ma, Nanyin Zhang

## Abstract

During anesthetic-induced unconsciousness (AIU), the brain undergoes a dramatic change in its capacity to exchange information between regions. However, the spatial distribution of information exchange loss/gain across the entire brain remains elusive. In the present study, we acquired and analyzed resting-state functional magnetic resonance imaging (rsfMRI) data in rats during wakefulness and graded levels of consciousness induced by incrementally increasing the concentration of isoflurane. We found that, regardless of spatial scale, the *absolute* functional connectivity (FC) change was significantly dependent on the FC strength at the awake state across all connections. This dependency became stronger at higher doses of isoflurane. In addition, the *relative* FC change (i.e. the FC change normalized to the corresponding FC strength at the awake state) exhibited a spatially homogenous reduction across the whole brain particularly after animals lost consciousness, indicating a globally uniform disruption of meaningful information exchange. To further support this notion, we showed that during unconsciousness, the entropy of rsfMRI signal increased to a value comparable to random noise while the mutual information decreased appreciably. Importantly, consistent results were obtained when unconsciousness was induced by dexmedetomidine, an anesthetic agent with a distinct molecular action than isoflurane. This result indicates that the observed global reduction in information exchange may be agent invariant. Taken together, these findings provide compelling neuroimaging evidence suggesting that the brain undergoes a widespread, uniform disruption in the exchange of meaningful information during AIU, and that this change may represent a common systems-level neural mechanism of AIU.

## Introduction

For over a century, the neural mechanisms of consciousness have been explored in the neuroscience field (Alkire et al. 2008). One common strategy to investigate such a phenomenon is the study of altered forms of consciousness as modulated in pharmacological, physiological or pathological conditions. In particular, <u>**a**</u>nesthetic-<u>**i**</u>nduced <u>**u**</u>nconsciousness (**AIU**) is an intriguing avenue for investigating consciousness given the convenience of experimental control and the essential role of anesthesia in modern medicine. AIU is a temporary and reversible loss of consciousness (LOC) produced by the administration of an anesthetic agent(s) and accompanied by analgesia, amnesia and immobility (Brown et al. 2010). Although the molecular actions of various anesthetic agents are fairly well understood, the systems-level neural mechanisms of AIU remain obscure.

Accumulating evidence suggests that AIU is a brain-network phenomenon. Anesthetics appear to suppress consciousness by disrupting information exchange across large-scale brain networks (Alkire et al. 2000; Tononi 2004; Mashour and Alkire 2013; Mashour 2014; Lee et al. 2013; Liang et al. 2012b; Boveroux et al. 2010; Martuzzi et al. 2010). This notion has been supported by a number of studies in both humans (White and Alkire 2003; Boveroux et al. 2010; Deshpande et al. 2010; Martuzzi et al. 2010) and animals (Vincent et al. 2007; Moeller et al. 2009; Wang et al. 2011; Liang et al. 2013), which have indicated that a disruption of functional connectivity (FC) specifically within the thalamocortical and frontoparietal networks is essential to AIU (Angel 1991; Velly et al. 2007; Lee et al. 2009; Breshears et al. 2010; Imas et al. 2005).

While these previous studies have highlighted the importance of thalamocortical and frontoparietal networks in AIU, whether and how the rest of the brain is involved during AIU remains unclear. The answer to this question will provide critical insight for clarifying whether the effects anesthetics are limited in specific neural networks (e.g. the thalamocortical network), or whether AIU leads to information exchange loss/gain across the whole brain. To address this issue, it is essential to elucidate changes across whole-brain networks during AIU. The importance of such investigation is also highlighted by the highly inter-connected characteristics of the brain organization, in which *global* coordination is critical for effective information exchange (Moon et al. 2015; Liang et al. 2012b). In addition, most anesthetic agents affect neurotransmitters throughout the entire brain (Franks 2008; Brown et al. 2010). Therefore, unraveling the effects of anesthesia on the whole-brain network can be critical for deciphering the systems-level neural mechanisms of AIU.

In the present study, we employed the technique of resting state functional magnetic resonance imaging (rsfMRI) to investigate information exchange loss/gain across the whole rat brain during wakefulness and graded levels of consciousness induced by increasing concentrations of isoflurane. rsfMRI uses the temporal correlation of low-frequency spontaneous fluctuations of the blood-oxygenation-level dependent (bold) signal as a measure of FC between brain regions (Biswal et al. 1995). Our data revealed that relative to the awake state, the FC change after LOC is reduced in a relatively uniform manner across the whole brain, which indicates that the disruption of information transfer is ubiquitous in the brain. In addition, this spatial pattern of information exchange loss was similar during LOC induced by a distinct anesthetic agent, dexmedetomidine. Collectively, these data suggest that the global disruption of information exchange may be an agent-invariant mechanism of AIU.

## Methods

### Animal Preparation

Thirty-seven adult male Long-Evans rats (300–500g) were housed and maintained on a 12hr light: 12hr dark schedule, and provided access to food and water *ad libitum*. Approval for the study was obtained from the Institutional Animal Care and Use Committee of the Pennsylvania State University. Animals were first acclimated to the scanning environment for 7 days to minimize stress and motion during imaging at the awake state (described in (Liang et al. 2011, 2012a, b, 2014; Liang et al. 2013; Liang et al. 2015a; Liang et al. 2015b; Zhang et al. 2010)). To do this, rats were restrained and placed in a mock MRI scanner where prerecorded MRI sounds were played. The exposure time in the mock scanner started from 15 minutes on day 1, and was incrementally increased by 15 minutes each day up to 60 minutes (days 4, 5, 6 and 7). This setup mimicked the scanning condition inside the magnet.

### Behavioral Assessment

The animal’s consciousness level was assessed using the behavioral test of loss of right reflex (LORR), which is a well-established index of unconsciousness in rodents. Researchers have demonstrated a strong correlation between the concentration of various anesthetic agents needed to produce the loss of voluntary movement in rodents and loss of consciousness in humans (unresponsiveness to verbal commands) over five orders of magnitude (Franks 2008). In the LORR procedure, each rat began in a restrained, awake state (i.e. 0% isoflurane) and was exposed to increasing doses of isoflurane through a nosecone. We measured the LORR for a single dose delivered to the rat exactly as it would be in the scanner (Figure 1): Each scan was 10 minutes with a 5-minute transition time between scans to allow the isoflurane concentration to reach a steady state. Thus, for example, to measure LORR at 1.5% isoflurane, the animal first received 0.5% isoflurane for 15 minutes, then 1.0% isoflurane for 15 minutes, and finally 1.5% isoflurane for 5 minutes. With the nosecone in place, the rat was then taken out of the restrainer and turned over to a supine position while the time it took to correct its position was recorded. If the rat did not correct its position within 60 seconds, it was deemed completely unconscious. This procedure mimicked anesthesia administration during the scanning process and allowed us to verify the animal’s consciousness level at each anesthetic dose administered during rsfMRI data acquisition.

**Figure 1:**
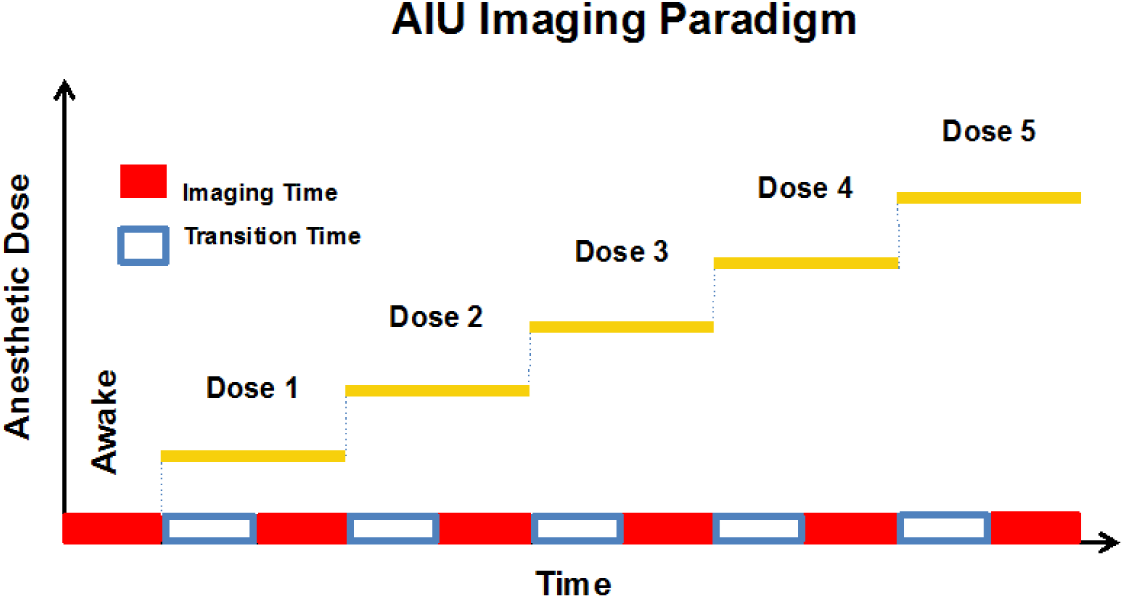
A schematic of the AIU imaging paradigm. Imaging began while animals were in a fully awake state, followed by five doses of increasing concentrations of isoflurane. For each dose, a 5-minute transition period was given before the rsfMRI scan started to allow the isoflurane concentration to reach steady state.

### MRI Experiments

Rats were briefly anesthetized with isoflurane while they were placed in a head restrainer with a built-in birdcage coil. Isoflurane was discontinued, but the nosecone remained fastened around the animal’s nose for the duration of the experiment. Imaging began 30 minutes after rats were placed in the scanner while animals were fully awake. Image acquisition was performed on a 7T scanner. A high-resolution T1 structural image was first acquired with the following parameters: repetition time (TR) = 2125 ms, echo time (TE) = 50 ms, matrix size = 256 × 256, field of view (FOV) = 3.2 cm × 3.2 cm, slice number = 20, slice thickness = 1 mm. rsfMRI data were then obtained during the awake state and graded levels of consciousness by systematically increasing the concentration of isoflurane from 0% (i.e. awake state) to 3% (0%, 0.5%, 1.0%, 1.5%, 2.0%, 3.0%). Isoflurane was administered via the nosecone. rsfMRI data were acquired using a single-shot gradient-echo, echo planar imaging pulse sequence with TR = 1000 ms, TE = 13.78 ms, flip angle = 80°, matrix size = 64 × 64, FOV = 3.2 × 3.2 cm, slice number = 20, 1mm thick slices (in-plane resolution = 500 m × 500 m). For each rsfMRI scan, 600 volumes were acquired in 10 minutes. Between each dose, 5 minutes were allowed to ensure the new dosage reached steady state (see Figure 1).

### Image Preprocessing

The first 10 volumes of each rsfMRI run were discarded to allow magnetization to reach steady state. The data was preprocessed with conventional procedures as previously described (Liang et al. 2014; Liang et al. 2015a; Liang et al. 2015b), including: registration to a segmented rat brain atlas, motion correction with SPM12, spatial smoothing (FWHM=1mm), regression of nuisance signals, as well as band-pass filtering (0.01-0.1Hz). 16 nuisance signals were regressed out, including the signals from whiter matter (WM), cerebrospinal fluid (CSF), 6 motion parameters and their derivatives. This regression strategy has been reported to minimize the impact of motion artifacts on rsfMRI data (Power et al. 2015). We chose not to regress out the global signal in order to avoid any spurious anticorrelations that may arise (Murphy et al. 2009).

### Data Analysis

Data were separately analyzed on a region-of-interest (ROI) and voxel basis. For the ROI-based analysis, we first parcellated the brain into 134 unilateral ROIs based on the anatomic definition of brain regions in Swanson Atlas (Swanson 2004). We derived FC by correlating the bold signal time series of each pair of ROIs. To ensure that our results were independent of the parcellation scheme and spatial scale, we also conducted the voxel-based analysis by calculating the FC between each pair of voxels within the brain. For each connection (between a pair of ROIs or between a pair of brain voxels), its absolute FC change was determined by subtracting the FC strength at the awake state from the corresponding FC strength at each anesthetized condition. In addition, the absolute FC change at each dose was divided by the corresponding FC strength (with sign) at the awake state (referred to as normalized FC change hereafter). Furthermore, to examine the change of information exchange for a local brain region (a voxel or ROI), we averaged the normalized FC change between that voxel (or ROI) and all other brain voxels (or ROIs) at each anesthetized condition (Olde Dubbelink et al. 2013). This averaged normalized FC change was used as the quantity to measure information exchange loss/gain at the voxel/ROI.

To test our hypothesis that spontaneous brain activity becomes more random during AIU as a result of meaningful information processing loss, we calculated the Shannon entropy of the bold time series for each ROI as well as the mutual information between each pair of ROIs at each condition. Entropy represents the expected uncertainty (or randomness) of the information source and mutual information is the reduction of this uncertainty. In the present study, entropy characterizes the information contained in the bold signal while mutual information represents the information exchanged between brain regions. To estimate the entropy distribution of random noise for the purpose of comparison, we simulated noise using a normal distribution and smoothed the noise using the same temporal filter used in data preprocessing. All data analyses were conducted using Mathematica software (Wolfram, Champaign, IL, USA).

## Results

In the current study, we examined the anesthetic-induced changes in information exchange across the whole brain as animals transitioned from an awake state to a unconscious state. rsfMRI data during wakefulness and at graded levels of consciousness produced by increasing concentrations of isoflurane (0.5%, 1.0%, 1.5%, 2.0% and 3.0%) were collected in rats.

### Animals lost consciousness at 1.5% isoflurane

To assess the animal’s consciousness level at each dose, we first measured the LORR outside of the scanner in a manner that directly mimicked the way the animal experienced the dose during rsfMRI scanning (Figure 1, also see **Methods** for details). We found that when 1.5% isoflurane was administered via a nosecone, 81% of animals were unable to correct from the supine to prone position. This percentage was significantly higher than when lower doses of isoflurane were administered (no rats lost righting reflex at lower doses, Figure 2, Chi-square test, χ^2^=22.82, p<2×10^−6^). Therefore, 1.5% was the dose to induce LOC when isoflurane was delivered through a nosecone. Notably, this dose was higher than typical isoflurane doses inducing LORR reported in the literature (0.7%-0.8%), in which animals were either intubated or were placed in an enclosed chamber (MacIver and Bland 2014; Hudetz et al. 2011). This difference is due to the different setup used in the present study (i.e. isoflurane was delivered via a nosecone), in which the actual concentration of isoflurane in the animal was lower than the concentration delivered due to air dilution around the nose cone.

**Figure 2:**
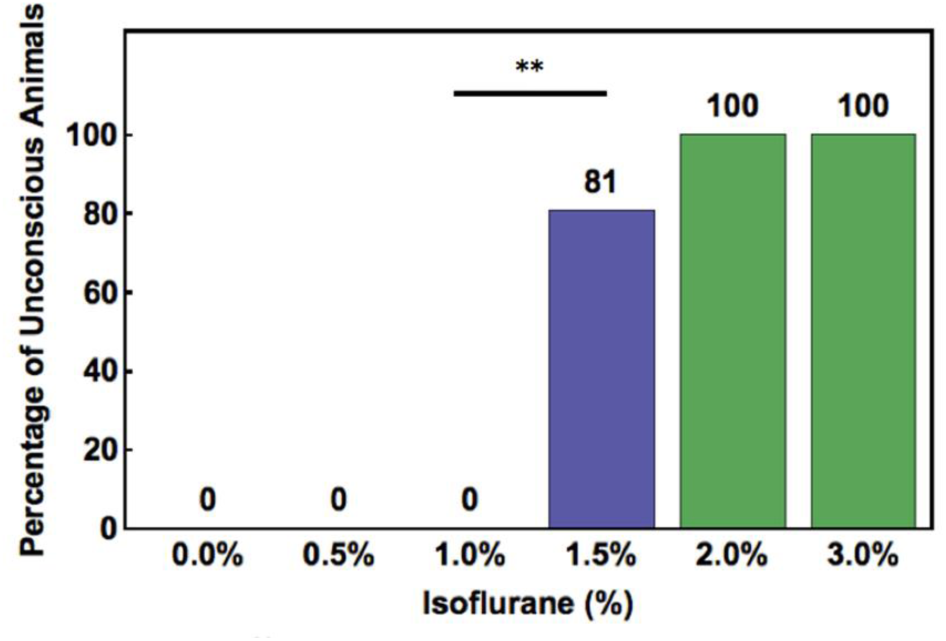
LORR test in isoflurane-anesthetized animals. The percentage of animals that were unable to correct from a supine to a prone position was measured at multiple concentrations of isoflurane: 0% (i.e. the awake state), 0.5%, 1.0%, 1.5%, 2.0% and 3.0%. This percentage significantly increased from 1.0% to 1.5% (Chi-square test, χ^2^=22.82, p<2×10^−6^), indicating that most animals lost consciousness at the isoflurane concentration of 1.5% (delivered through a nosecone).

### FC was significantly disrupted in the thalamocortical and frontoparietal networks

Numerous literature studies (White and Alkire 2003; Boveroux et al. 2010; Deshpande et al. 2010; Martuzzi et al. 2010; Vincent et al. 2007; Moeller et al. 2009; Wang et al. 2011; Liang et al. 2013; Scott et al. 2014; Hudetz et al. 2015; Liu et al. 2011, 2013b) have reported that AIU is associated with a disruption of FC within the thalamocortical and frontoparietal networks. To replicate this finding, we examined the absolute FC changes between ROIs in the thalamus, primary sensory-motor, parietal and prefrontal cortices (see Table 1 for the list of ROIs) between the awake state and an unconscious state (3% isoflurane) using the conventional subtraction method. Figure 3 shows connections with significantly decreased (blue lines) and increased (red lines) FC within the thalamocortical and frontoparietal networks (in the axial and coronal views, two sample t-test, p < 0.05, False Discovery Rate corrected). The data indicate that a large number of connections within these two networks were compromised during AIU. This result confirms the findings reported in the literature.

**Figure 3:**
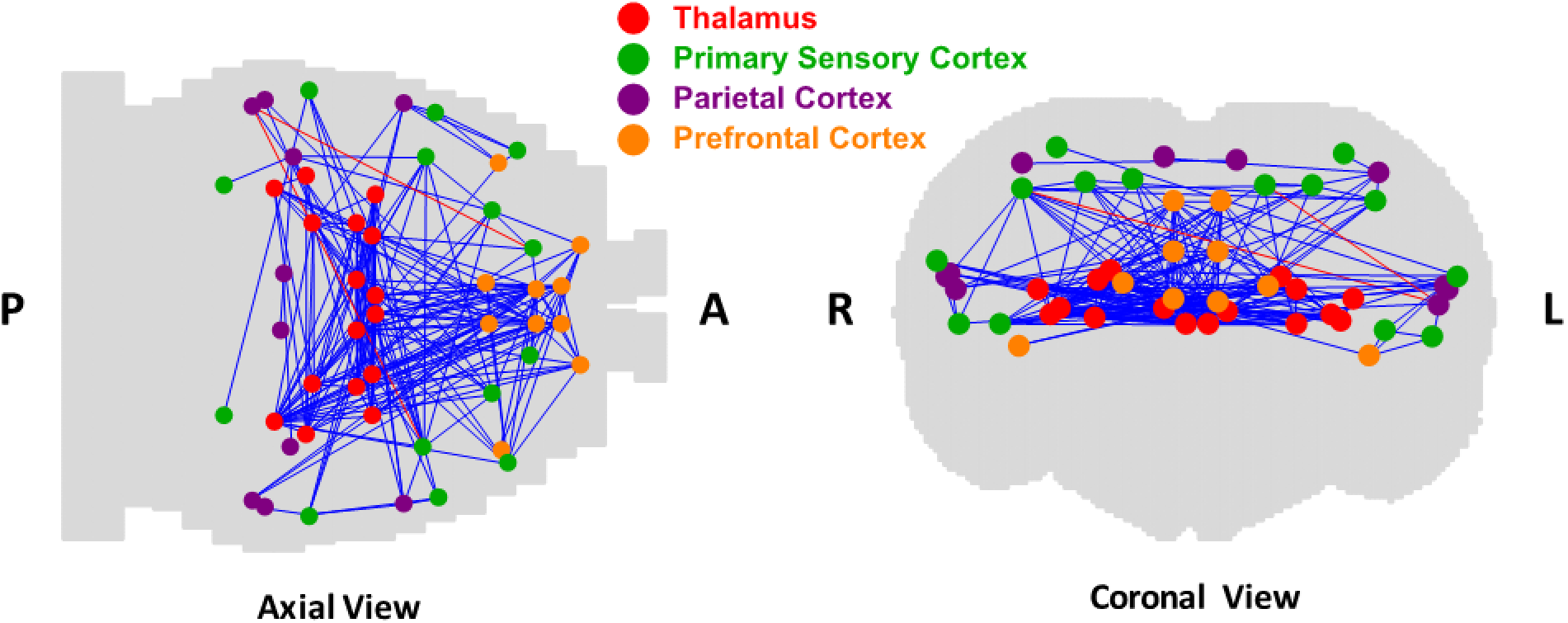
Significant absolute FC changes within the thalamocortical and frontoparietal networks at an unconscious state. Significantly decreased (blue lines) and increased (red lines) FC between ROIs (listed in Table S1) in the thalamus, primary sensory, parietal, and prefrontal cortices (color-coded) from the awake state to 3.0% isoflurane displayed in the axial and coronal views of the rat brain (t-test, p < 0.05, False Discovery Rate corrected). P: posterior; A: anterior; R: right; L: left.

**Table 1:** Brain regions of interest in the thalamus, primary sensory-motor, parietal and prefrontal cortices.

| **Thalamus** | **Primary sensory cortex** | **Parietal cortex** | **Prefrontal cortex** |
| --- | --- | --- | --- |
| Anterior nuclei, dorsal thalamus | Auditory areas | Ectorhinal area | Agranular insular area |
| Lateral geniculate complex | Gustatory area | Parietal region | Anterior cingulate area |
| Lateral nuclei, dorsal thalamus | Primary somatomotor area | Retrosplenial area | Infralimbic area |
| Medial geniculate complex | Primary somatosensory area | Supplemental somatosensory area | Orbital area |
| Medial nuclei, dorsal thalamus | Secondary somatomotor area | Ventral temporal association areas | Prelimbic area |
| Midline group, dorsal thalamus | Visceral area |  |  |
| Reticular nucleus thalamus | Visual areas |  |  |
| Ventral nuclei, dorsal thalamus |  |  |  |

### The absolute FC change during AIU was significantly dependent on the FC strength at the awake state across all connections in the brain

We further examined anesthetic-induced FC changes across the whole brain. The brain was parcellated into 134 unilateral ROIs. For each dose, we calculated the absolute FC change (i.e. ΔFC) between every two ROIs relative to the awake state (134 unilateral ROIs and 8911 connections in total). Intriguingly, this *ΔFC highly depended on the corresponding FC strength at the awake state across all connections in the brain* (Figure 4A, r= −0.41, −0.49, −0.62, −0.60, −0.81, for isoflurane concentrations of 0.5%, 1.0%, 1.5%, 2.0% and 3.0%, respectively, p<10^−200^ for all concentrations). In addition, this dependency became stronger at higher doses of isoflurane, reflected by increasingly negative correlations between ΔFC and the FC strength at the awake state when the isoflurane dose increased. The difference in these correlations was statistically significant across doses (one-way ANOVA, F(4,120) = 5.95, p =0.0002, Figure 4C). Furthermore, this dependency existed regardless of the spatial scale as evidenced by the consistency of the results obtained for all connections between every pair of brain voxels (Figure 4B, ~6000 voxels and ~18 million connections, r= −0.60, −0.64, −0.71, −0.72 and −0.79 for isoflurane concentrations of 0.5%, 1.0%, 1.5%, 2.0% and 3.0%, respectively, p<10^−200^ for all concentrations). The difference in these correlations was also statistically significant across doses for the voxel-wise analysis (one-way ANOVA, F(4,120) = 3.32, p =0.01, Figure 4D).

**Figure 4:**
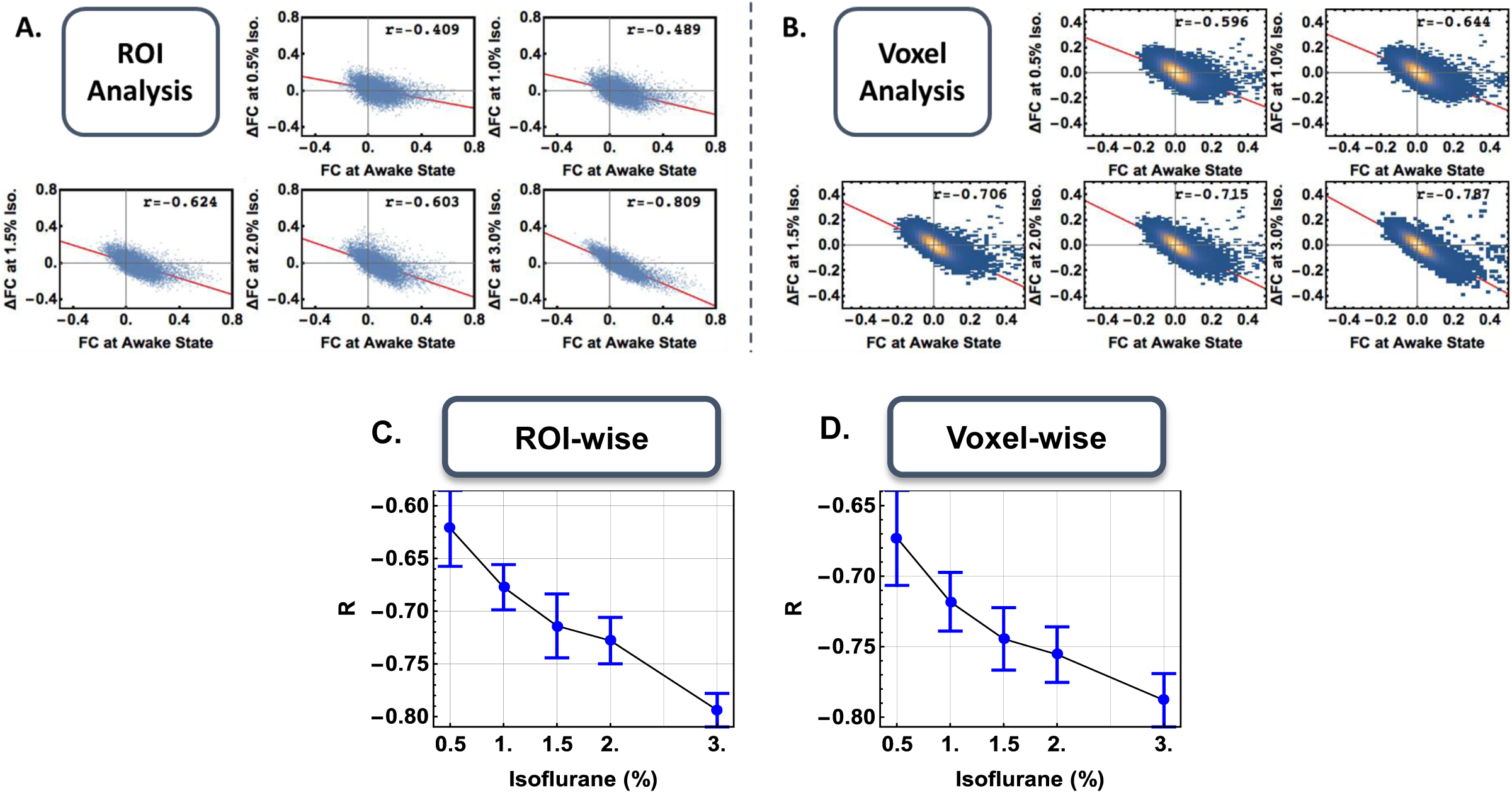
The absolute changes in FC are dependent on the FC strength at the awake state. (A, B) Scatter plots of (A) ROI-based and (B) voxel-based analyses show that the dependency between the absolute FC changes at anesthetized states and the FC strength at the awake state becomes stronger at higher doses of isoflurane. (C, D) Correlation coefficient (r) between the absolute FC changes and the FC strength at the awake state as a function of isoflurane concentration in both ROI- (C) and voxel-wise (D) analyses. The difference in these correlations was statistically significant across doses for both the ROI (one-way ANOVA, F(4,120)=5.95, p=0.0002) and the voxel-wise (one-way ANOVA, F(4,120)=3.32, p=0.01) analyses. Bars: SEM.

### The brain undergoes widespread, uniform disruption in information exchange during isoflurane-induced unconsciousness

To investigate the information exchange loss/gain at a local brain region during AIU, we first normalized the absolute FC change of each connection (i.e. ΔFC) at each anesthetized condition to the corresponding FC strength at the awake state (referred to as *normalized FC change*). For each brain voxel (or ROI), we then averaged its *normalized FC changes* with all other brain voxels (or ROIs), and this quantity was used as a measure of information exchange loss/gain for this voxel (or ROI) (Olde Dubbelink et al. 2013). Figure 5 shows voxel-wise histograms (A) and spatial distributions (B) of the averaged *normalized FC change* for all isoflurane doses. Strikingly, all voxels exhibited negative *normalized FC change* for all doses, indicating an exclusive reduction of information exchange under anesthesia. In addition, the averaged *normalized FC change* became spatially homogeneous at unconscious states (i.e. isoflurane dose ≥ 1.5%). These results collectively suggest a spatially exclusive and relatively uniform information exchange loss widely across the whole brain during AIU.

**Figure 5:**
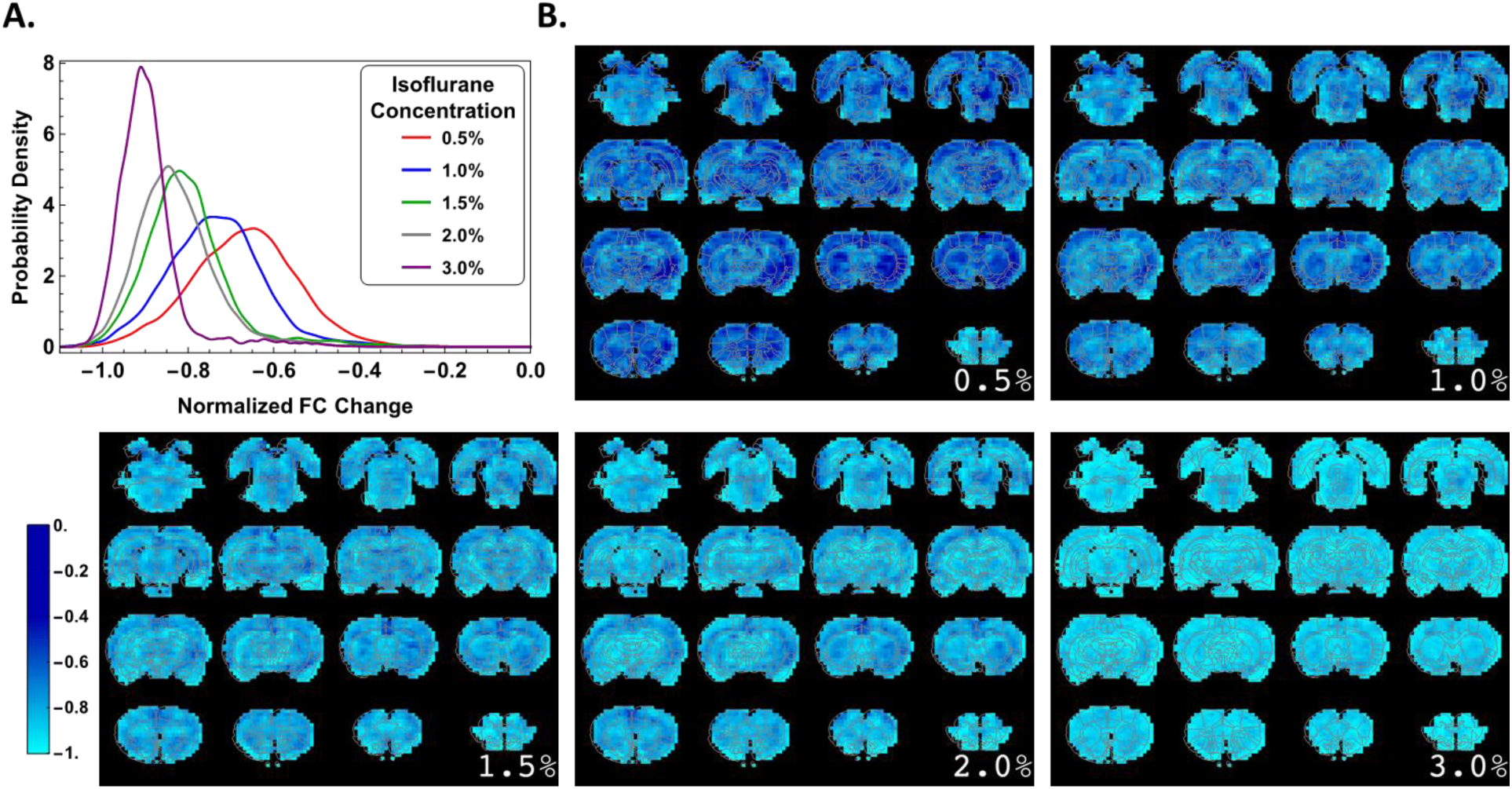
Distributions and voxel-wise maps of the averaged normalized FC changes at each dose of isoflurane. (A) Histograms show that all voxels exhibited negative *normalized FC change* for all doses, indicating an exclusive reduction of information exchange under anesthesia. (B) Maps reveal a spatially uniform reduction in normalized FC once animals lost consciousness (isoflurane concentration ≥ 1.5%).

Figure 5A also demonstrated that, relative to the awake state, the overall FC strength across the brain was reduced by ~80% (i.e. normalized FC change < −0.8) after the animal lost consciousness (i.e. isoflurane dose ≥ 1.5%). If the vast majority of meaningful information processing throughout the brain was lost during AIU, it can be predicted that the randomness of spontaneously fluctuating neural activity and rsfMRI signal should significantly increase (Barttfeld et al. 2015). To test this hypothesis, we calculated Shannon entropy and mutual information of the bold signal at each condition. Here, entropy, which provides a measure of randomness in the bold signal, characterizes the amount of information that the signal contains, and mutual information describes the amount of information that is exchanged. Figure 6A shows that the entropy of the bold time series increased with increasing isoflurane concentration and remained stable beyond the point the animal first lost consciousness (1.5% isoflurane). This entropy change was statistically significant across doses (one-way ANOVA, F(5,144) = 4.99, p = 0.0003). Notably, the entropy values during unconsciousness approach that of random noise (average entropy value of simulated random noise = 2.01). Conversely, mutual information between brain regions gradually declined as the level of consciousness decreased (Figure 6B). This change in mutual information was also statistically significant across doses (one-way ANOVA, F(5,144)=6.20, p=0.0003). Taken together, these data again support that meaningful information exchange was significantly disrupted across the whole brain during isoflurane-induced unconsciousness.

**Figure 6:**
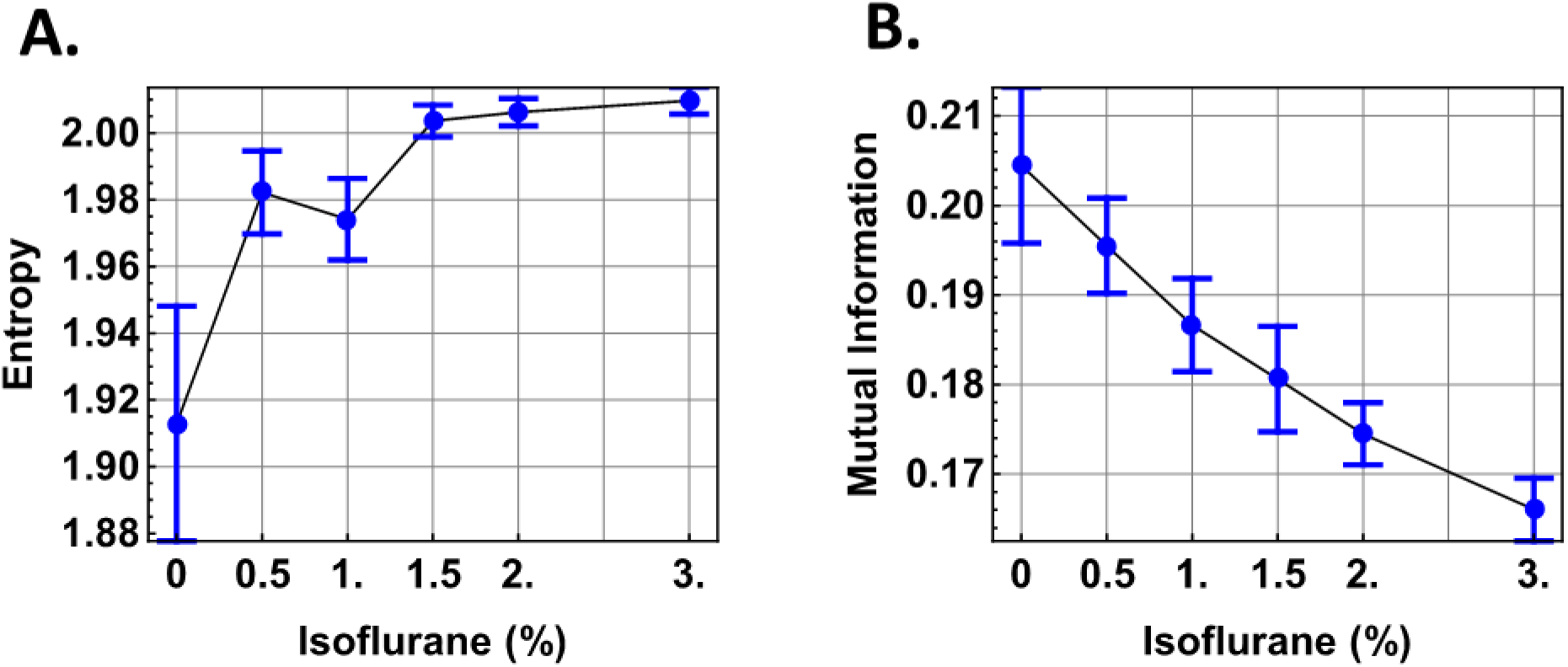
Entropy and mutual information at each isoflurane dose. (A) The entropy of the bold signal increases to a value corresponding to random noise (~2.0), and (B) mutual information appreciably decreases as anesthetic depth increases. Bars: SEM across rats.

### Global reduction in information exchange is not agent specific

To examine whether the global pattern of information exchange disruption during isoflurane-induced unconsciousness is agent-dependent, and also to rule out the possibility that the changes we observed result from the vascular effects of isoflurane (Ori et al. 1986; Lenz et al. 1998; Maekawa et al. 1986; Alkire et al. 1997), we further investigated FC changes during dexmedetomidine-induced unconsciousness. Dexmedetomidine is an alpha-2-adrenoceptor agonist with virtually no vascular effect, and its molecular action is distinct from isoflurane (Nelson et al. 2003). We acquired rsfMRI scans at both the awake and unconscious states induced by subcutaneous injection of a bolus of 0.05 mg/kg of dexmedetomidine (Angel 1993; Nelson et al. 2003; Pawela et al. 2008; Zhao et al. 2008; Weber et al. 2006), followed by a continuous infusion of dexmedetomidine (0.1 mg/kg/h) initiated 15 minutes after the bolus injection to maintain sedation (Angel 1993; Nelson et al. 2003; Pawela et al. 2008; Zhao et al. 2008; Weber et al. 2006). The dose selected was strong enough to abolish the righting reflex in all animals (n=6). rsfMRI data were analyzed in the same way as above.

Similar to isoflurane-induced unconsciousness, we observed a strong dependency between the absolute FC changes at the dexmedetomidine-induced unconscious state and the FC strength at the awake state across all connections (Figure 7A, ROI-wise analysis, r = −0.74, p < 10^−200^; Figure 7B, voxel-wise analysis: r = −0.72, p < 10^−200^). Consistent results between ROI- and voxel-wise analyses indicate that information exchange loss was independent of the spatial scale under dexmedetomidine. Also similar to isoflurane-induced unconscious states, the normalized FC changes were drastically reduced by ~85% during dexmedetomidine-induced unconsciousness (Figure 7C), and the spatial distribution of this information exchange loss was homogenous across the brain (Figure 7D). Moreover, like isoflurane-induced unconsciousness, we observed an increase in the entropy (Figure 7E) and a decrease in mutual information (Figure 7F) during dexmedetomidine-induced unconsciousness. Collectively, these data suggest that the global disruption of information exchange is not agent specific and may be an agent-invariant mechanism of AIU.

**Figure 7:**
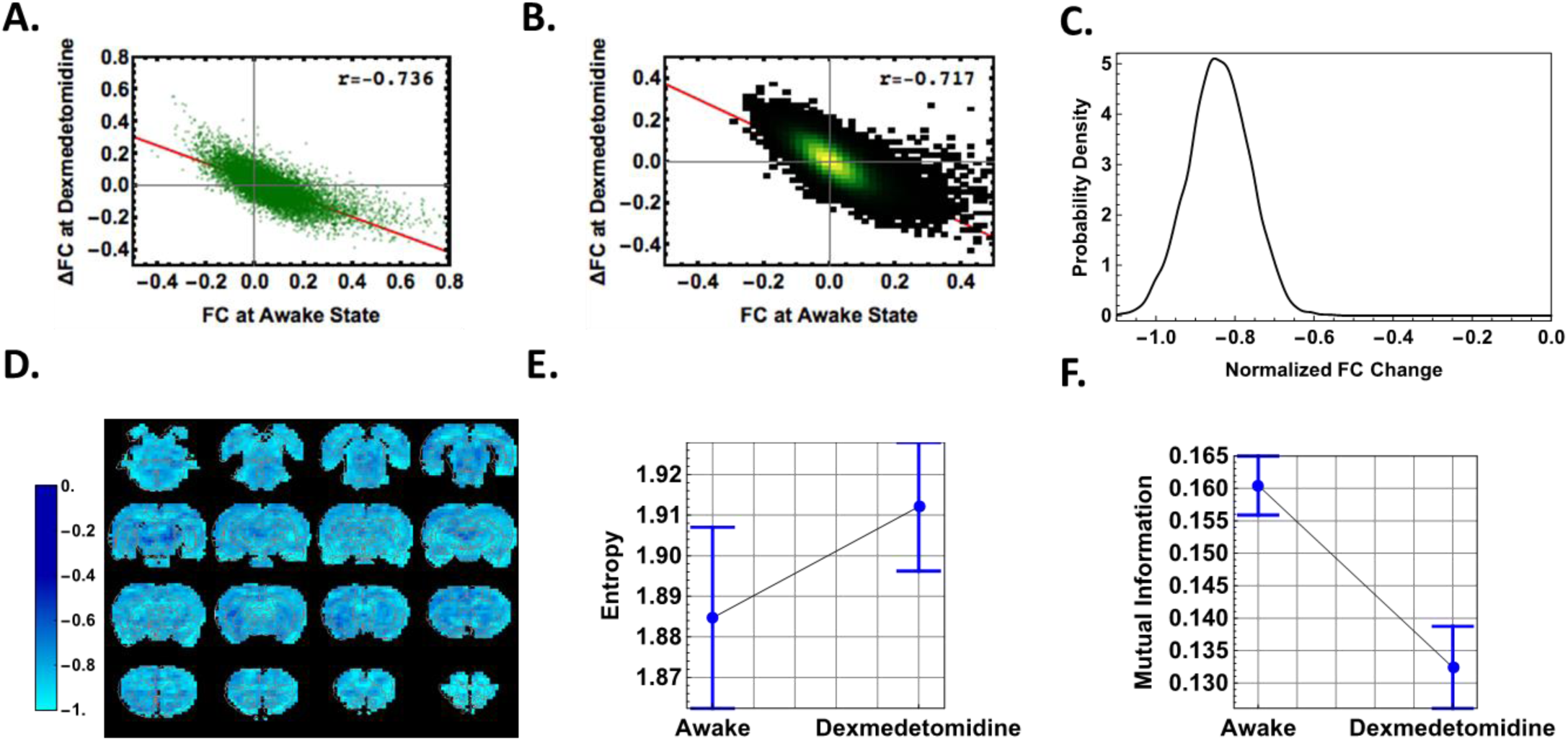
FC changes during dexmedetomidine-induced unconsciousness. (A, B) The absolute FC changes at the anesthetized state are dependent on the FC strength at the awake state for both ROI- (A) and voxel-based (B) analyses. (C, D) The distribution (C) and voxel-wise map (D) of the normalized FC change during dexmedetomidine-induced unconsciousness. (E, F) The entropy (E) and mutual information (F) of the bold signal at the awake state and during dexmedetomidine. Bars: SEM.

## Discussion

In the present study, we investigated the spatial pattern of anesthetic-induced FC changes across the whole brain in rats. We found that the absolute FC changes at anesthetized states depended on the FC strength at the awake state. This dependency became stronger at higher doses of isoflurane, and persisted regardless of the spatial scale. To further investigate the regional distribution of information exchange loss/gain during AIU, for each voxel (or ROI) we averaged its normalized FC changes with all other voxels (or ROIs). Our analysis indicates that the brain exhibited exclusive reduction in this averaged normalized FC during unconsciousness, reflecting the brain’s diminished capacity to exchange meaningful information between regions. To further support this notion, we demonstrate that after LOC, the entropy of the bold signal approached a level comparable to random noise and the mutual information showed a marked reduction. Taken together, these results indicate that during AIU, information exchange is significantly disrupted across the whole brain.

### Anesthesia has a global, rather than local, impact on the brain

Since the brain is a highly inter-connected system in which *global* coordination is critical for effective information processing (Moon et al. 2015), and also considering that most anesthetic agents affect neurotransmitters ubiquitously (Franks 2008), it is reasonable to hypothesize that anesthesia has a global impact on brain functioning. To test this hypothesis, a technique that allows whole-brain networks to be studied at different consciousness levels is needed. rsfMRI is ideal for this purpose given its features of global field of view, superb spatial resolution and high sensitivity to dynamic changes. In addition, rsfMRI allows us to quantify functional communication that occurs when two brain regions generate synchronized neural activity, and thus can provide a measure of information exchange at different states. By using this technique, we demonstrate that information exchange loss is exclusive and relatively uniform across the brain during AIU. These results confirm that anesthesia has a global, rather than local, impact on the brain.

### Global information exchange loss is agent independent

Our data also provide important evidence that the global reduction of information exchange during AIU might be independent of the anesthetic agent used, as we show consistent findings in rats anesthetized by dexmedetomidine—an anesthetic agent with a drastically different molecular mechanism from isoflurane. Isoflurane is a halogenated ether that affects both neural and vascular substrates (Angel 1993), whereas dexmedetomidine is an alpha-2-adrenoceptor agonist with virtually no vascular effect. Similar patterns of information exchange loss between dexmedetomidine- and isoflurane-induced unconsciousness can rule out the possibility that the FC changes we observed originated from the vasodilating effect of isoflurane (Lenz et al. 1998). More importantly, these result suggest that global information exchange loss may be a common phenomenon across various anesthetic agents, or even, a general mechanism of unconsciousness. This notion is supported by a recent study demonstrating disruption of corticocortical information transfer during ketamine anesthesia (Schroeder et al. 2016), and report of breakdown of FC during propofol-induced LOC (Boly et al. 2012; Liu et al. 2013a). Future studies can extend this analysis to more anesthetic agents and to alternative forms of unconsciousness, such as sleep or coma.

### Global reduction of information exchange is associated with increased randomness of spontaneous brain activity

If information exchange indeed diminishes across the whole brain during AIU, one would predict that meaningful information processing would be significantly compromised, giving rise to more random spontaneous neural activity. This notion is supported by our findings that, as anesthetic depth increases, the entropy of the rsfMRI signal increased to a value comparable to random noise (~2.0), and mutual information decreased appreciably. These results are consistent with an independent study reporting that, at a higher isoflurane concentration (2.9%), the temporal scaling characteristics of rsfMRI signal, measured by Hurst exponent (H), approached those of Gaussian white noise (H = 0.5), in contrast to lower isoflurane concentrations (0.5%-1%, H>0.5) (Wang et al. 2011). They also agree with a recent study in which Barttfeld and colleagues (2015) demonstrated that FC at unconscious states arises from “a semi-random circulation of spontaneous neural activity along fixed anatomical routes”, producing a single brain connectivity pattern dictated by the structure of anatomical connectivity (Barttfeld et al. 2015). These results collectively suggest that global reduction of information exchange is associated with increased randomness of spontaneous brain activity.

### Normalized FC changes can reveal subtle FC differences

The relatively uniform reduction of FC across the whole brain during unconsciousness appears to contradict many neuroimaging studies that highlight specific brain regions/circuits that are particularly sensitive to anesthesia (e.g. thalamocortical and frontoparietal networks). However, this apparent contradiction is most likely attributed to the difference in data analysis methods and the interpretation of results. Previous neuroimaging studies have relied on the subtraction method that statistically compares absolute FC changes between the awake and anesthetized states (Barttfeld et al. 2015) or between different anesthetic depths (Hutchison et al. 2014; Vincent et al. 2007). As a result, these studies tend to highlight regions/networks that exhibit large FC changes. Indeed, by using the conventional subtraction method our data showed significantly reduced FC in the thalamocortical and frontoparietal networks as well (Fig. 3). Intriguingly, we also found that, in general connections with stronger FC in the awake state tend to exhibit larger absolute FC changes during AIU (Fig. 4). Therefore, to reveal more subtle FC changes, and also to account for the influence of the FC strength in the awake state, we normalized the absolute FC changes at each anesthetic depth to the FC strength at the awake state. Consequently, this analysis provides a global, holistic perspective of whole-brain connectivity changes during AIU. To demonstrate that the conventional subtraction method would reveal apparently different results, we identified connections with significant absolute FC changes (red dots in Figure 8, p < 0.005, uncorrected). It is clear from this figure that with the use of the subtraction method, a specific set of connections was highlighted, while the global-scale proportional relationship revealed in the present study would no longer be as pronounced.

**Figure 8:**
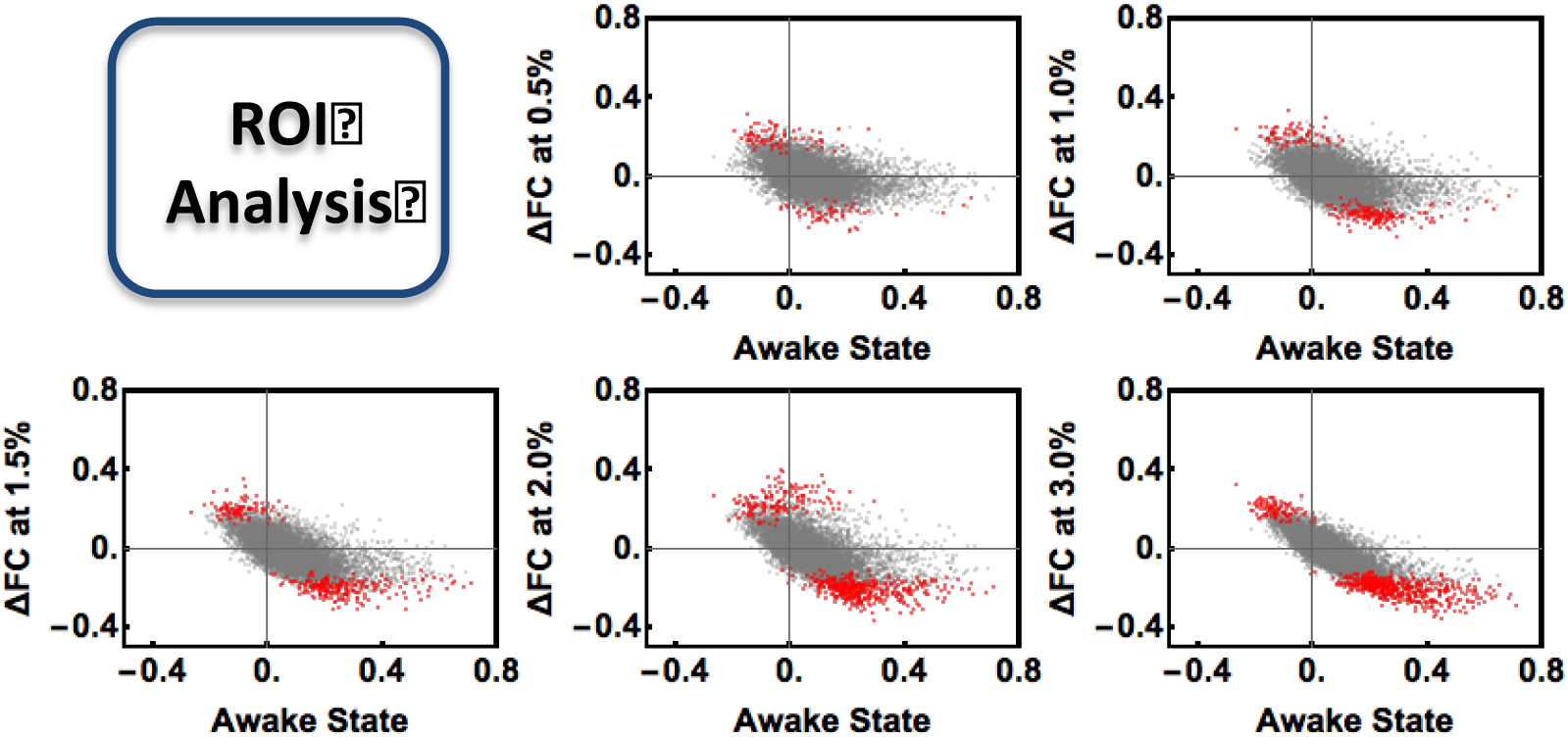
Scatter plots illustrating the relationship between the absolute FC changes (between pairs of ROIs) and the strength of the FC at the awake state for each dose of isoflurane. The red points represent the connections that exhibited significant absolute FC changes using the conventional subtraction method (two-sample t-test, p<0.005, uncorrected).

### Advantages and Limitations

One major advantage of the present study is the utilization of the awake animal imaging approach (Liang et al. 2011; Zhang et al. 2010). Animal fMRI experiments typically rely on anesthesia to immobilize animals, which confounds the effects of the anesthetic agents being studied. Consequently, without the ‘ground truth’ of brain activity/connectivity at the awake state, it is virtually impossible to parcel out the specific effects of an anesthetic agent on global brain networks, and thus is difficult to fundamentally decipher systems-level mechanisms underpinning AIU. This obstacle has been overcome in the present study, in which rsfMRI data of awake animals were collected and used as the reference point to determine the whole-brain FC changes during various unconscious conditions.

One potential limitation is the higher motion level in the rsfMRI data at the awake state compared to anesthetized states (averaged volume-to-volume displacement = 0.148 mm for the awake state, and 0.066 mm for all anesthetized states). However, the disparity in motion between different consciousness levels cannot explain the relatively uniform FC reduction we observed during AIU. First, it is unlikely that larger motion at the awake state can result in uniformly stronger FC (Van Dijk et al. 2012). Second, in our data preprocessing we used a 16-parameter nuisance regression approach, which has been shown to be very effective for removing motion-related artifacts in FC calculations (Power et al. 2015). More importantly, we identified a subset of data (n = 7) that exhibited minimal movement during the awake state (averaged volume-to-volume displacement = 0.068mm at the awake state), and this motion level was similar to that at anesthetized states (one-way ANOVA: F(4,36) = 2.08, p=0.1). Figure 9 shows that the pattern of information exchange loss obtained from this subset of data is almost identical to that from the full dataset, suggesting that our results cannot be attributed to difference in motion.

**Figure 9:**
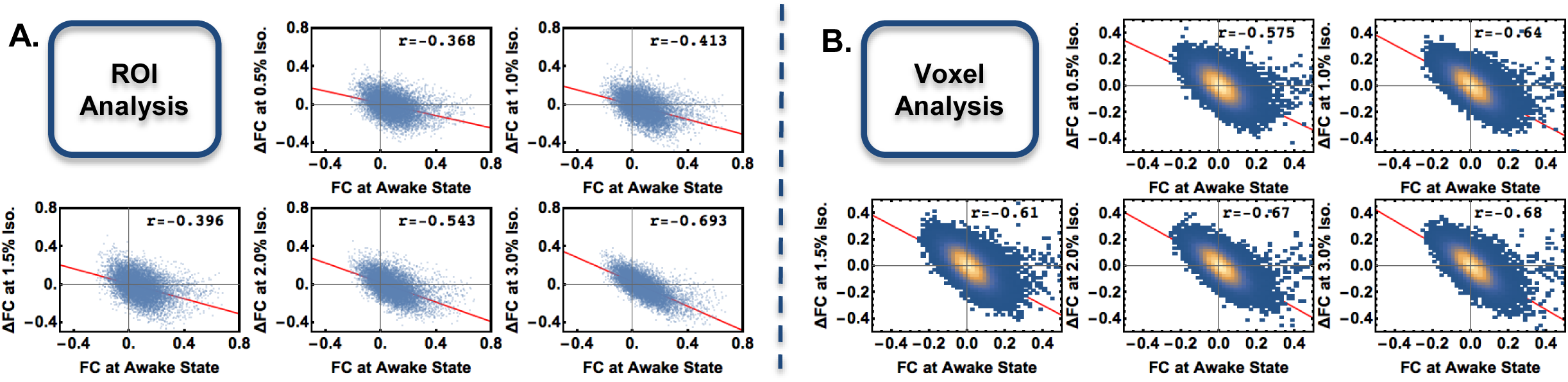
(A) ROI- and (B) voxel-based analyses of a subset of data that exhibited minimal movement during the awake state. In this subset of data, the motion levels of the awake and anesthetized states were comparable (one-way ANOVA: F(4,36)=2.08, p=0.1). Almost identical results obtained in the subset suggest that our findings are not attributed to differences in the motion level across different consciousness levels.

### Summary

Overall, the present study provides compelling neuroimaging evidence showing a global reduction of FC during AIU, which reflects the disruption of meaningful information exchange across the whole brain. This notion is bolstered by the increase in entropy of the bold signal and a marked reduction in mutual information during unconsciousness. Because this pattern was evident during unconsciousness induced by two distinct anesthetic agents, we suspect that this mechanism may be agent invariant. These findings allude to a common systems-level mechanism of AIU in which the information exchange capacity is affected in all brain regions/networks to a similar degree.

The uniform reduction of FC associated with anesthetized states might serve as a biomarker for unconsciousness that could be utilized in a clinical setting. Such a biomarker would effectively reduce anesthesia-related complications during surgical procedures. Broadly, this research yields novel insight that contributes to our scientific understanding of consciousness with the potential of uncovering fundamental features of this phenomenon. Such information would be especially valuable to individuals suffering from disorders of consciousness that are non-communicative and non-responsive (e.g. Lock-in Syndrome), as its translation into a clinical context would facilitate both diagnosis and prognosis of these conditions.

## Acknowledgement

We would like to thank Dr. Pablo Perez for his technical support, and Ms Lilith Antinori for editing the manuscript. The work was supported by the National Institutes of Health, Grant Numbers R01MH098003 (PI: Nanyin Zhang, PhD) from the National Institute of Mental Health and R01NS085200 (PI: Nanyin Zhang, PhD) from the National Institute of Neurological Disorders and Stroke.

